# Global trends in human birth and death seasonality reveal climate-related shifts in reproductive timing

**DOI:** 10.64898/2026.01.20.700719

**Authors:** Chinmay Lalgudi, Neema Sakariya, James E. Ferrell, Shixuan Liu

## Abstract

Despite growing evidence that climate change affects public health, it remains unclear whether recent global warming has directly altered fundamental aspects of human biology at large scales. Here, we analyzed long-term trends in human birth and death seasonality, using demographic data from >100 regions worldwide. We found that both births and deaths followed remarkably persistent seasonal cycles. Births peaked through spring to fall, whereas deaths peaked in mid-winter, especially among older individuals. Across latitude, death seasonality diminished near the equator; birth seasonality remained strong across all latitudes but peaked progressively later at lower latitudes. Strikingly, birth seasonality in high-latitude regions showed synchronized phase delays in recent decades, with peak timing delayed by 2–3 months during the 1970s–2000s, converging toward lower-latitude timing and stabilizing subsequently, while death seasonality remained stable. An unbiased analysis of key socioeconomic and climatic variables revealed that temperature was the predominant predictor of birth-peak timing. Estimated conception peaks shifted away from summer amid rising heatwaves and stabilized in cooler fall/winter months, consistent with known adverse effects of heat on reproduction. Together, these results suggest that recent climate warming has reshaped the seasonal timing of human reproduction.

## Introduction

In nature, many organisms align key life cycle events with predictable seasonal changes. Seasonal reproduction is widespread across many taxa.^1^ Seasonal breeders in temperate regions typically give rise to offspring in spring or summer, when conditions are most favorable for survival. However, because gestation or incubation lengths vary across species, mating occurs at different times to ensure that births coincide with favorable seasons. For example, small animals with short gestation periods generally mate in early spring, whereas large mammals with longer gestation tend to mate in the preceding fall or winter. This timing involves well-coordinated physiological control, but the molecular basis of this regulation is only partly understood.^1–4^ Mortality also shows seasonal clustering in some species,^5,6^ but these patterns are less well documented, and it remains unknown whether any intrinsic physiological regulation is involved beyond environmental constraints.

Unlike classical division into seasonal versus non-seasonal breeders, seasonal reproduction spans a continuum, from strict seasonal breeders with narrow reproductive windows to species that reproduce across much or all of the year, yet still show clear seasonal patterns in births.^7–9^ While humans are classically viewed as continuous breeders, they nonetheless exhibit clear seasonal variations in birth rates. Seasonal patterns in human births have been recognized for centuries, and seasonal patterns in mortality have also been long documented.^10–12^ However, reported seasonality patterns can differ substantially across regions, creating long-standing inconsistencies that have complicated a unifying understanding. More recent studies have often focused on specific countries or regions, with an emphasis on disease associations and other epidemiological implications, and have occasionally reported weakening or temporal changes in seasonality within specific populations.^13–24^ Whether these variations reflect a coherent underlying pattern or arise from disparate, population-specific influences has yet to be resolved.

Seasonal regulation in humans is likely shaped by multiple environmental cues, such as photoperiod, temperature, and resource availability.^25–27^ Recently, climate change has emerged as an important modifier of seasonal environmental conditions,^28^ with well-documented effects on public health, including mortality during extreme climate events, heat-related morbidity, and disruptions to food and disease systems.^29–32^ Yet, it remains unclear whether recent climate change has directly altered human physiology at large scales.

Here, we assembled a global dataset of birth and death seasonality based on data from 159 countries and regions across all major inhabited latitudinal zones. This systematic analysis found remarkably persistent seasonal cycles of both births and deaths worldwide, and identified distinct latitudinal patterns in their seasonality that help reconcile long-standing geographical inconsistencies. We further examined how seasonality patterns have changed over time, using data extending back to the 1840s. Strikingly, all analyzed high-latitude regions experienced pronounced shifts in their peak birth timing between about 1970 and 2010; births previously peaked in the spring to early summer, and have now shifted to the summer and fall, approaching the timing seen in low latitude regions. In contrast, deaths have peaked consistently in mid-winter across latitudes during this period. An unbiased analysis of 13 socioeconomic and climate variables identified temperature as the predominant predictor of birth season, explaining both the geographic gradient in birth peak phase and its temporal shift over recent decades. Warmer regions and years exhibited later birth peaks, even after adjusting for latitude. Indeed, estimated conception peaks at high latitudes shifted away from summer amid rising heatwaves and stabilized in the cooler fall and winter months. In contrast, at lower latitudes, conception peaks remained stable in the coolest winter months over the same period. These patterns are consistent with the well-established heat sensitivity of male fertility, and suggest that recent climate warming, especially the increasing frequency of summer heat extremes, may have already reshaped the seasonal timing of human reproduction.

## Results

### A global multi-decade dataset of monthly birth and death rates

To examine seasonal patterns of births and deaths in humans, we analyzed demographic datasets of live births and deaths across 159 countries and regions worldwide (Table S1). We focused on datasets reported in the United Nations (UN) database,^33^ which compiles standardized international data at monthly resolution. We also analyzed additional datasets that complemented the main UN data by including longer temporal coverage and, in some cases, breakdowns of deaths by age, sex, and underlying cause of death (see below).

Figure 1a-c shows world maps of the 159 countries/regions included in the analysis, which span all six inhabited continents and numerous island territories across the major oceans. Notable absences from the database included mainland China, India, Indonesia, and much of Africa. Temporal coverage varied across countries/regions. UN birth data go back to 1967, while death data are available from 1980 onward. Both extend through the 2020s, although occasional gaps (i.e., years with no or incomplete entries) are common in individual countries/regions. The latitude of the most populous city was used as a proxy for each country/region. We excluded countries/regions with fewer than 3 years of data or insufficient numbers of recorded births or deaths to ensure robust analyses (see Methods). This resulted in 124 countries/regions for subsequent analysis (122 for birth, 95 for death), with an average of 30.6 ± 15.6 (mean ± SD) years of birth data and 29.1 ± 11.6 years of death data per country/region.

**Figure 1.**
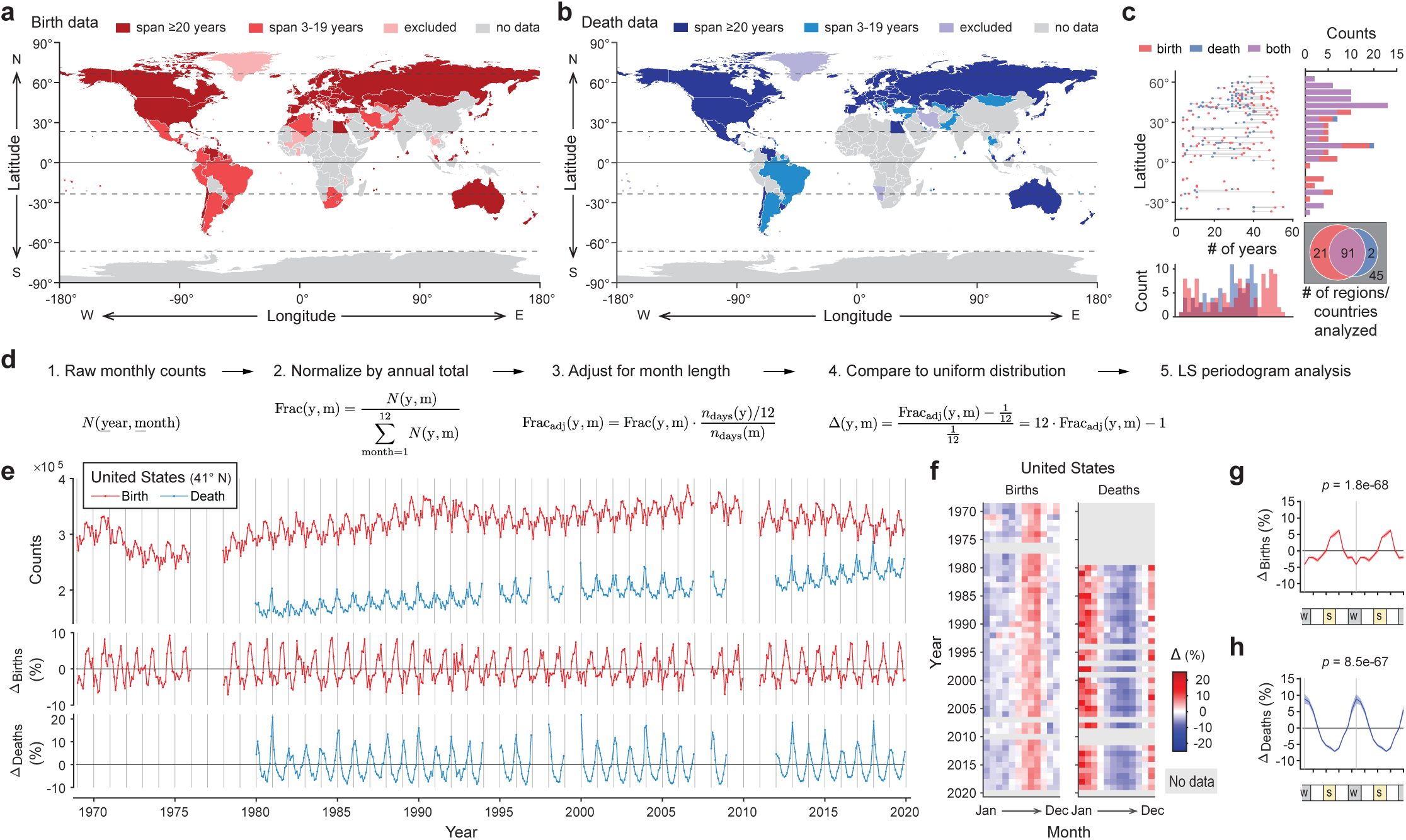
A global multi-decade dataset of monthly birth and death rates. **a-c**, World maps and temporal coverage of birth (a) and death (b) data, and their summary statistics (c). The latitude of the most populous city was used as a proxy for the country/region. Some regions were excluded from further analysis due to quality control reasons, as described in Methods. **d**, Schematic of the data normalization and seasonality analysis pipeline. **e**, Time-series of raw birth and death counts (top) and percent deviations in the rates of birth (Δbirths, middle) and death (Δdeaths, bottom) in the United States (US). **f**, Heat-maps showing Δbirths (left) and Δdeaths (right) in the US across months and years. **g-h**, Average annual patterns and 95% confidence intervals of Δbirths (g) and Δdeaths (h) in the US, averaged across years. Patterns are double plotted for two cycles for visualization purposes. Vertical gray lines in (e, g, h) indicate January.

To calculate monthly birth and death rates, raw counts were normalized by annual total and adjusted for the number of days in each month (Fig. 1d). We then calculated percent deviations of birth and death rates relative to a uniform distribution, referred to as Δ_births_ and Δ_deaths_, respectively. Seasonality was assessed using Lomb-Scargle (LS) periodograms applied to each time series, and average annual birth and death patterns were derived by averaging across years for each country/region (Fig. S1a-b).

The United States (US) is shown as an example (Fig. 1e-h). Raw counts of births and deaths exhibited clear seasonal patterns (Fig. 1e, top), which became smoother after normalization (Fig. 1e, middle and bottom). Births peaked consistently around August and September, and deaths in January (Fig. 1e-f). Both the birth and death rates displayed highly significant 12-month periodicity (birth, *p* = 1.8e-68; death, *p* = 8.5e-67), with an average amplitude (half peak-to-trough difference) of 5.8 ± 0.8% for births and 9.8 ± 2.3% for deaths (Fig. 1g-h). Thus, the US exhibited robust, periodic changes in both birth and death rates, and the amplitude and phase of the seasonal patterns varied little over the last three to five decades.

### Human births and deaths are seasonal worldwide, but show different patterns across latitude

Using the same approaches, we examined global patterns of human births and deaths across all 124 countries/regions that passed quality control (Table S1, Fig. 1a-b). To avoid potential confounding effects of the COVID-19 pandemic, we excluded death data from 2020 onward in this analysis, reducing the death data to 93 countries/regions (see Methods).

This global analysis revealed widespread seasonality in human births (Fig. 2a-b) and deaths (Fig. 2e-f). LS periodograms detected significant (*p* ≤ 0.05) 12-month periodicity in 90% (110/122) of the countries/regions analyzed for births and 85% (79/93) for deaths (Fig. 2a, e). Most of them (68%, 83/122 for births; 67%, 62/93 for deaths) showed highly persistent seasonality, with 12-month periodicity *p*-values < 1e-10. To compare patterns between Northern and Southern Hemispheres, we shifted the Southern Hemisphere data by 6 months to make the seasons align. This alignment revealed consistent trends of births and deaths by season in both hemispheres, with clear symmetry across the equator (Fig. 2b, f).

**Figure 2.**
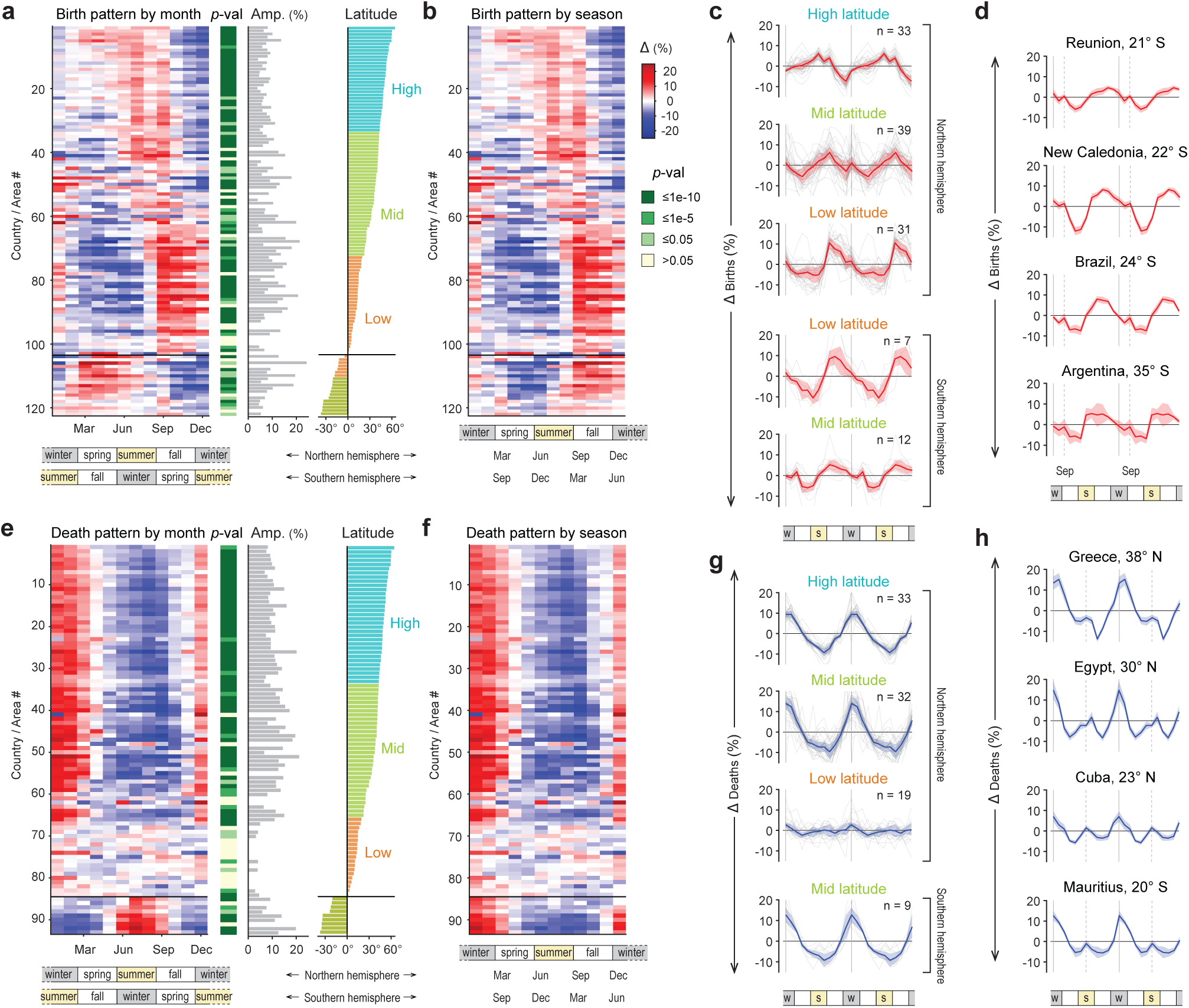
Human births and deaths are seasonal worldwide, but show different patterns across latitude. **a, e**, Heatmaps showing average annual patterns of Δbirths (a) and Δdeaths (e) for different countries/regions (rows), ordered from northmost to southmost latitudes. Panels adjacent to each heatmap show the *p*-values for 12-month periodicity, average cycle amplitude (Amp.) for regions with *p* < 0.05, and latitude of corresponding country/region. Amplitude is defined as half the peak-to-trough-difference in Δbirths and Δdeaths. **b, f**, Heatmap of Δbirths (b) and Δdeaths (f) as in (a) and (e), but with Southern Hemisphere data shifted 6 months to align countries/regions by season rather than calendar month. **c, g**, Average annual patterns of Δbirths (c) and Δdeaths (g) across latitudinal zones, calculated by averaging countries/regions within each zone. **d**, Average annual patterns of Δbirths for selected countries in the southern mid-latitudes, showing minor September birth peaks that may reflect holiday-related conceptions. **h**, Average annual patterns of Δdeaths for selected countries in northern or southern mid-latitudes, showing secondary mid-summer death peaks that may reflect heat-wave-related mortality.

Birth peak timing showed an interesting dependency on latitude (Fig. 2a-c, Fig. S1c). At higher latitudes, births peaked earlier in spring to summer, whereas toward lower latitudes, the birth peaks gradually shifted later into late summer and fall. Notably, regions in the low latitudes and tropics showed pronounced birth seasonality, with amplitudes often greater than those observed at higher latitudes (Fig. 2a-c, Fig. S1c). In addition to the primary peak, a minor secondary birth peak was observed in the southern mid-latitudes (Fig. 2d), which is confined to September and likely reflects New Year holiday conceptions. However, this secondary peak had a much smaller amplitude and prominence than the main seasonal cycle, suggesting that conceptions associated with major social and cultural events around the New Year contribute only marginally to overall birth seasonality. Similarly, the timing of the largest holiday in each country/region did not correlate to its birth peak month (Fig. S1d).

Death peaks exhibited geographically more consistent patterns than those of births, with death rates peaking consistently in mid-winter (Fig. 2e-g, Fig. S1c). A latitudinal pattern was also evident, but differed from that of births. Seasonal variation in deaths was minimal near the tropics but became evident at mid- and high-latitudes. Interestingly, the amplitude of death seasonality did not increase gradually with latitude. Instead, pronounced seasonality appeared abruptly upon reaching the mid-latitudes, where seasonal cycle amplitudes were often as high as, or slightly higher than, those at higher latitudes (Fig. 2e-g, Fig. S1c). We also observed a secondary death peak in mid-summer in several mid-latitude countries (Fig. 2h), likely reflecting increased mortality associated with heatwaves.

### Death seasonality strengthens with age, and remains stable despite flu season shifts

For a small subset of countries, mortality records from national statistical agencies were stratified by finer demographic groups. The United Kingdom, for example, reported weekly deaths across seven age groups, ranging from age 0 (<1 year) to 85+ years. Older populations contributed a larger share of total deaths and exhibited notable seasonality, with consistent peaks in mid-winter (Fig. 3a-b). Significance and amplitude of death seasonality tended to increase with age (Fig. 3c, Fig. S2a). Among the oldest age group (age 85+), death rates at the winter peaks were about 40% higher than those at the summer troughs. There was little seasonal variation in the death patterns of the youngest age group (<1 year). The group of 1-14 year-olds showed a high seasonal amplitude (Fig. 3c) but the variability in the data was very high (Fig. S2a). No significant sex differences were observed (Fig. S2a). Other countries, including the Netherlands, Switzerland, and New Zealand showed similar trends, with death seasonality increasing with age (Fig. S2b-d).

**Figure 3.**
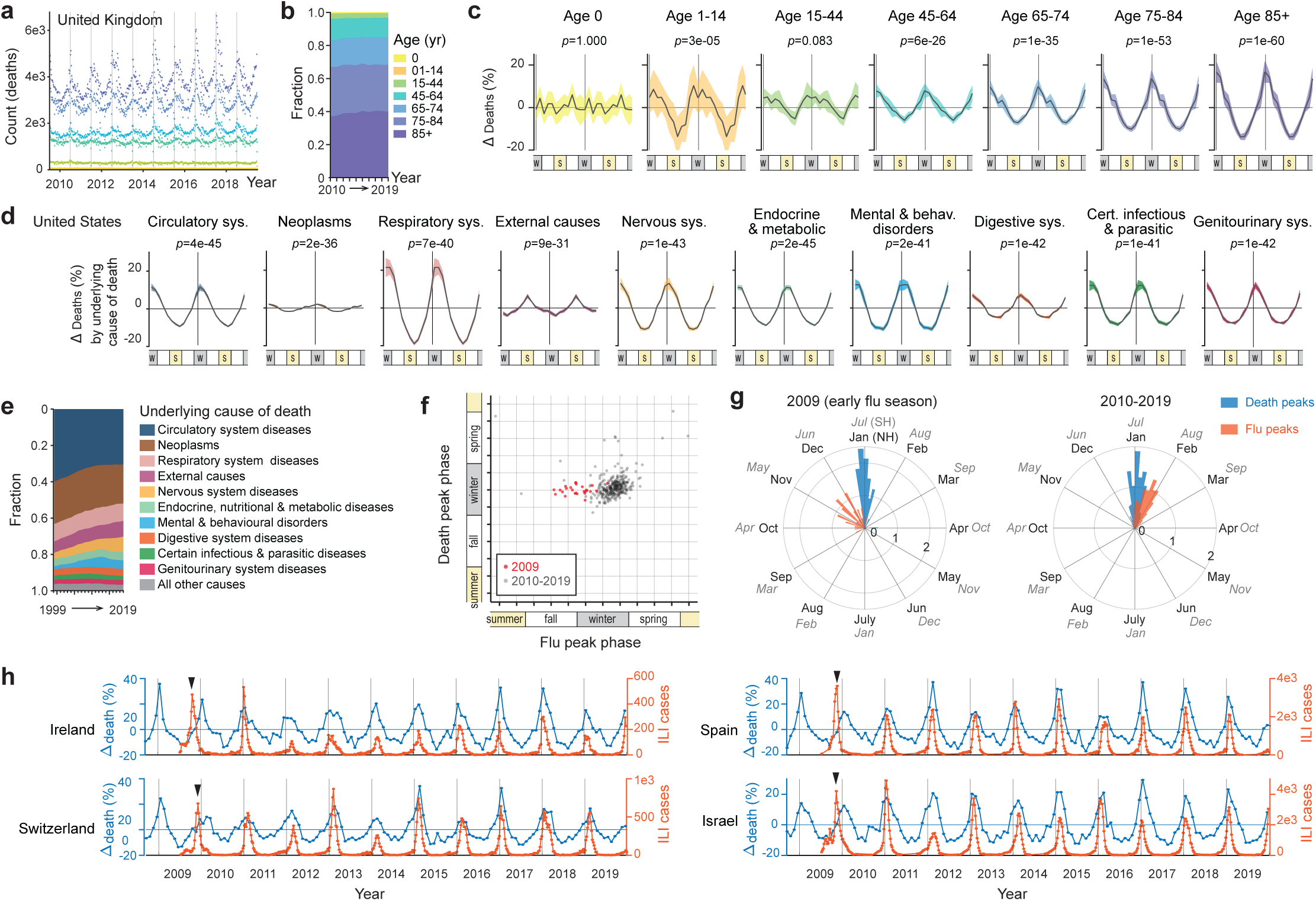
Death seasonality strengthens with age, and does not change in accordance with flu season shifts. **a-b**, Weekly death counts (a) and fractions (b) by age groups in the United Kingdom (UK) from 2010 to 2019. **c**, Average annual patterns of Δdeaths in the UK for different age groups, calculated from the data in (a-b). Curves and shaded areas show median and 95% confidence intervals. Patterns are double plotted for two cycles for visualization purposes. Vertical gray lines mark the beginning of the year. Note the strengthening of death seasonality over age from middle age to the elderly. See also Fig. S2. **d-e**, Average annual patterns of Δdeaths (d) and fractions (e) in the US separated into the top ten leading causes of death, formatted as in (c). Among the nine leading disease causes of death, most exhibit clear seasonal patterns with persistent winter peaks, except for neoplasms, which show little to no seasonality. Deaths associated with external causes (e.g., accidents) are weakly seasonal and peak in summer. **f**, Scatter plot comparing the timing of flu peaks (based on influenza-like illness (ILI) cases) versus that of death peaks, across years and countries/regions. **g**, Polar histograms comparing the timing of flu peaks and death peaks in the 2009-2010 flu season (left) and in subsequent years (2010-2019, right). The radial axis shows probability density. Inner month labels denote Northern Hemisphere (NH); outer month labels denote Southern Hemisphere (SH). **h**, Δdeaths and ILI cases over time in example countries. Gray vertical lines indicate the start of the year. Arrowheads mark the unusually early flu peak in 2009-2010. Note despite this advance, death peaks remained at their typical timing and did not advance accordingly.

A wide range of diseases, from seasonal affective disorder and viral infections to cardiovascular diseases, have long been recognized to exhibit persistent seasonal patterns.^18,19,34–41^ We therefore examined death seasonality by underlying cause of death in the United States, for which detailed data were available^42^ (Fig. 3d-e, Fig. S3). The top ten leading causes of death accounted for over 95% of total mortality, nine of which were disease-related (Fig. 3e). Although neoplasia, the second leading cause of death, showed little or no seasonality, the other eight disease categories all showed clear seasonal patterns with pronounced winter peaks (Fig. 3d, Fig. S3a). These included conditions related to the cardiovascular, respiratory, nervous, digestive, endocrine/metabolic, and genitourinary systems, as well as certain infectious/parasitic diseases. Deaths from external causes (e.g., accidents) showed only weak seasonality, and peaked instead in the summer (Fig. 3d, Fig. S3a). Death due to respiratory-system diseases showed the strongest seasonality, with the winter peaks about 50% higher than the summer troughs. A similar pattern was observed in the UK, where deaths were categorized into respiratory disease-related versus all other causes (Fig. S3b).

The winter influenza season is often assumed to underlie the winter peaks in mortality. Is death seasonality a direct consequence of the flu season? To address this question, we examined influenza surveillance data assembled at the World Health Organization’s Global Influenza Programme.^43^ This dataset reports weekly influenza-like illness (ILI) cases for individual countries/regions, beginning in the 2009-2010 flu season. We compared the time of ILI peaks to that of the death peaks across years and countries/regions. Consistent with previous reports,^44^ the 2009-2010 flu season was exceptionally early, with cases peaking between late October and December in many regions, well in advance of the typical January flu peaks (Fig. 3f-h). This natural experiment provided an opportunity to assess how the timing of flu seasons affects winter mortality. Interestingly, death peaks in the 2009-2010 cycle were not advanced and still occurred around January (Fig. 3f-h), suggesting that winter mortality is not simply driven by the timing of the flu season.

### Birth peaks shifted to later months at high-latitudes in recent decades

Having established the geographical and latitudinal patterns, we next analyzed temporal patterns of birth and death seasonality over the past decades to examine whether the amplitude or phase of seasonal peaks changed across years. To ensure sufficient temporal coverage and robustness, we restricted the analysis to countries/regions with at least 20 years of data in the UN database and highly significant 12-month periodicity (*p* < 1e-10). This yielded 73 countries/regions analyzed for birth and 56 for death, with average coverage of 40.2 ± 8.8 years (mean ± SD, birth) and 36.4 ± 5.3 years (death) of data per country/region (Table S1). We then calculated the seasonal cycle amplitude and peak phase for each country/region and year. Amplitude was defined as the half difference between the annual maximum and minimum monthly values. To obtain a robust estimate of peak phase, we calculated the weighted mean birth month (i.e. center of mass) across all 12 months using circular statistics (see Methods).

The amplitude of both birth and death seasonality appeared largely stable over the past four to five decades (Fig. S4a, c). This is surprising given the technological and socioeconomic transformations and increasing urbanization during this period, which have shifted much of the global population, especially in developing regions,^45^ into urban lifestyles that buffer exposure to seasonal variations in food availability and other environmental conditions.

Strikingly, we observed phase delays of birth peaks across high latitude regions (Fig. 4a-d, Fig. S4b). In Denmark, for example, birth peaks occurred in spring (around April) in the early 1970s, but gradually shifted later over the following decades to summer (July to August) in recent years (Fig. 4a). A similar phase delay was observed across all high latitude countries/regions analyzed (Fig. 4c-d), spanning most of Europe as well as parts of North and South America, including Canada and Chile (Fig. 4a-b). In contrast, lower-latitude areas such as Japan, Israel, Cuba, and Singapore, showed a largely stable birth peak phase in fall over the same period (Fig. 4a, c-d). High- and mid/low-latitude regions differed markedly in their temporal patterns (Fig. 4c-d, Fig. S4b). Within the high latitude regions, we also observed latitudinal variation in the phase delay (Fig. 4c). Regions closer to the poles initially exhibited earlier peaks in spring that shifted later and stabilized in early-to-mid summer, while slightly lower-latitude areas began with slightly later peaks (e.g., late spring to early summer) and stabilized in late summer to early fall. This resulted in a convergence of birth peak timing across latitudes, making high-latitude regions more similar to mid/low-latitudes (Fig. 4c-d). While a latitudinal gradient in peak timing remains, the overall variation across latitudes has become smaller than in the past.

**Figure 4.**
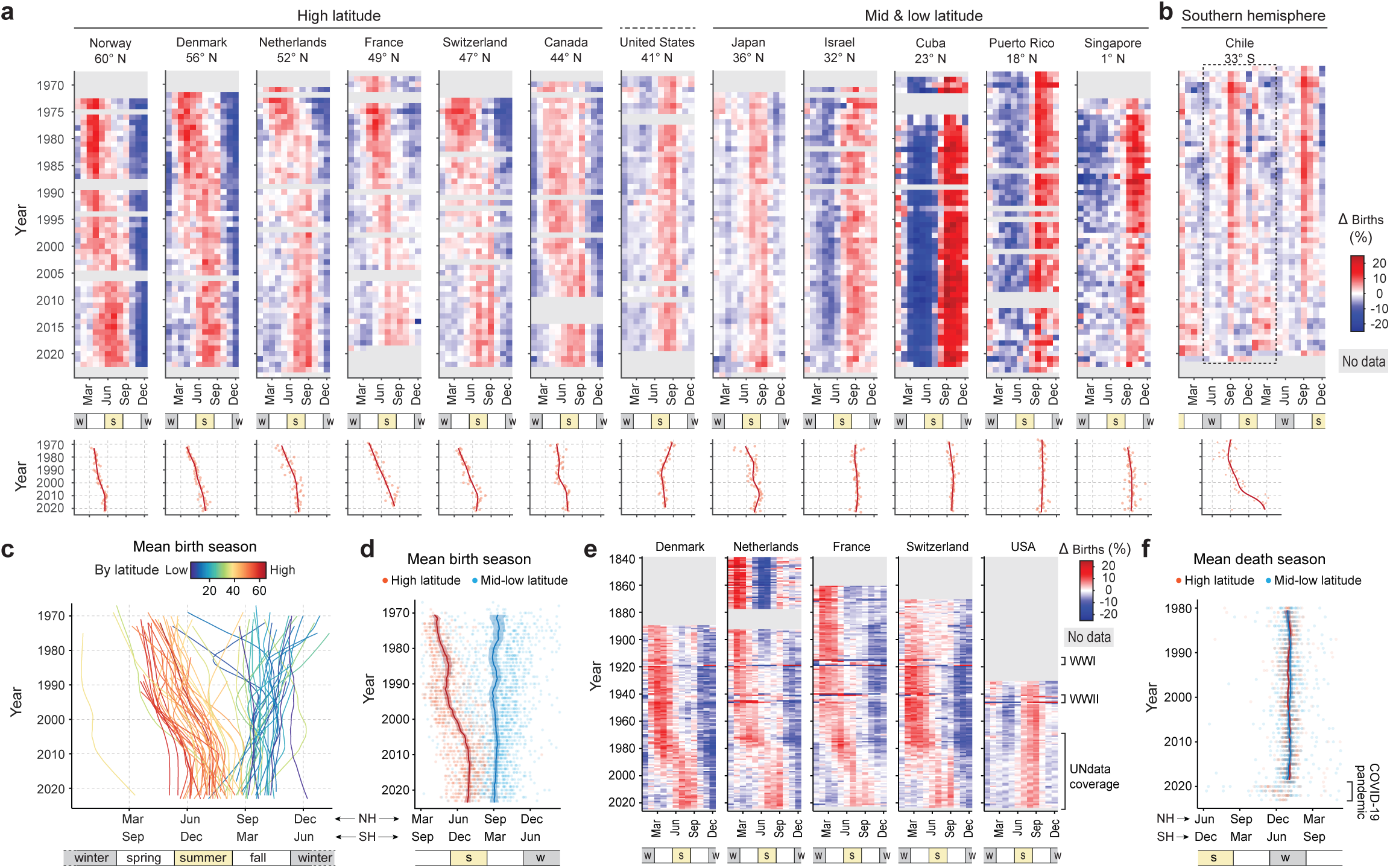
Birth peaks have shifted to later months at high-latitudes in recent decades. **a**, Heatmaps of Δbirths (top) for selected high- and mid/low-latitude countries, along with their calculated yearly birth peak phases and nonparametric regression curves (bottom). Note the gradual phase delay in birth peaks at high-latitudes in recent decades, shifting from spring/early summer toward mid/late summer and fall. **b**, Heatmap of Δbirths (top) and yearly birth peak phases (bottom) for a Southern Hemisphere country, Chile, also showing a gradual phase delay in birth peaks from spring toward summer/fall. **c**, Regression of yearly birth peak phases over time as in (a), overlaid across different countries/regions (curves) and colored by absolute latitude. Note the phase delays in high latitude regions and the relatively stable birth peak timing in mid/low latitudes. The leftmost curve likely reflects known reporting artifacts thus was excluded from subsequent analyses (see Methods). **d**, Birth peak phase for each country/region and year (dots), with nonparametric regression over time performed separately with high-latitude (> 43°) and mid/low-latitude (≤ 43°) data points. **e**, Heatmap of Δbirths for countries in the Human Fertility Database with longitudinal records extending back to the 1840s. Note the phase shift in high latitude countries between the 1970s and early 2000s, contrasted with relatively stable birth peak timing before and after this interval. Also note the brief irregularities during World War I (WWI) and World War II (WWII). **f**, Death peak phase for individual countries/regions and years (dots), with nonparametric regression over time performed separately with high-latitude (> 43°) and mid/low-latitude (≤ 43°) data points. Death peaks remained stable in midwinter across space and time, with noticeable disruptions only during the recent COVID-19 pandemic. NH: Northern Hemisphere; SH: Southern Hemisphere.

Birth peaks at high latitudes shifted at an average rate of 0.08 ± 0.03 (mean ± SD) months per year (i.e., 2.3 months over three decades) between the 1970s and early 2000s and appeared to have stabilized around 2000-2010 (Fig. 4c-d). However, this phase delay was already evident in the earliest UN records from around 1970. This raises the question of when the phase delay first began. To address this, we analyzed additional datasets from the Human Fertility Database,^46^ which largely overlap with the UN database but, for a few countries, provide records extending much further back, with the longest extending to the 1840s. Analysis of these data showed that most of the phase shift occurred between approximately the 1970s and 2000s, with relatively stable birth peaks before and after this interval (Fig. 4e). Interestingly, these data also captured the periods of the two World Wars, which caused short-term irregularities in birth patterns in these countries. However, birth seasonality quickly recovered after each war, returning toward pre-war patterns (Fig. 4e).

In contrast to birth peak patterns, death peaks remained remarkably stable over time, with consistent peaks in mid-winter across different latitudes since the beginning of the UN records in the early 1980s (Fig. 4f, Fig. S4d). The only notable deviations occurred during the recent COVID-19 pandemic, which caused occasional irregularities in death seasonality patterns (Fig. 4f, Fig. S4d). Together, these findings highlight a stark contrast between the seasonality of births and deaths. While birth peaks varied across latitude and shifted substantially over recent decades, death peaks remained consistent across space and time, suggesting regulation by different underlying mechanisms.

### Photoperiod and artificial-light exposure do not explain the temporal changes in birth peak timing

Photoperiod is an influential and well-characterized environmental cue regulating seasonal rhythms across a wide range of species.^1–3,47^ Indeed, we found a clear negative correlation (r = −0.84) between the natural annual photoperiod amplitude of a country/region and its birth peak phase (Fig. 5a), suggesting that photoperiod could explain the latitudinal gradient of birth peak timing. However, the natural photoperiod cycle itself is effectively constant over time with negligible changes on the scale of a few decades, and thus cannot account for the temporal shifts in birth peaks.

**Figure 5.**
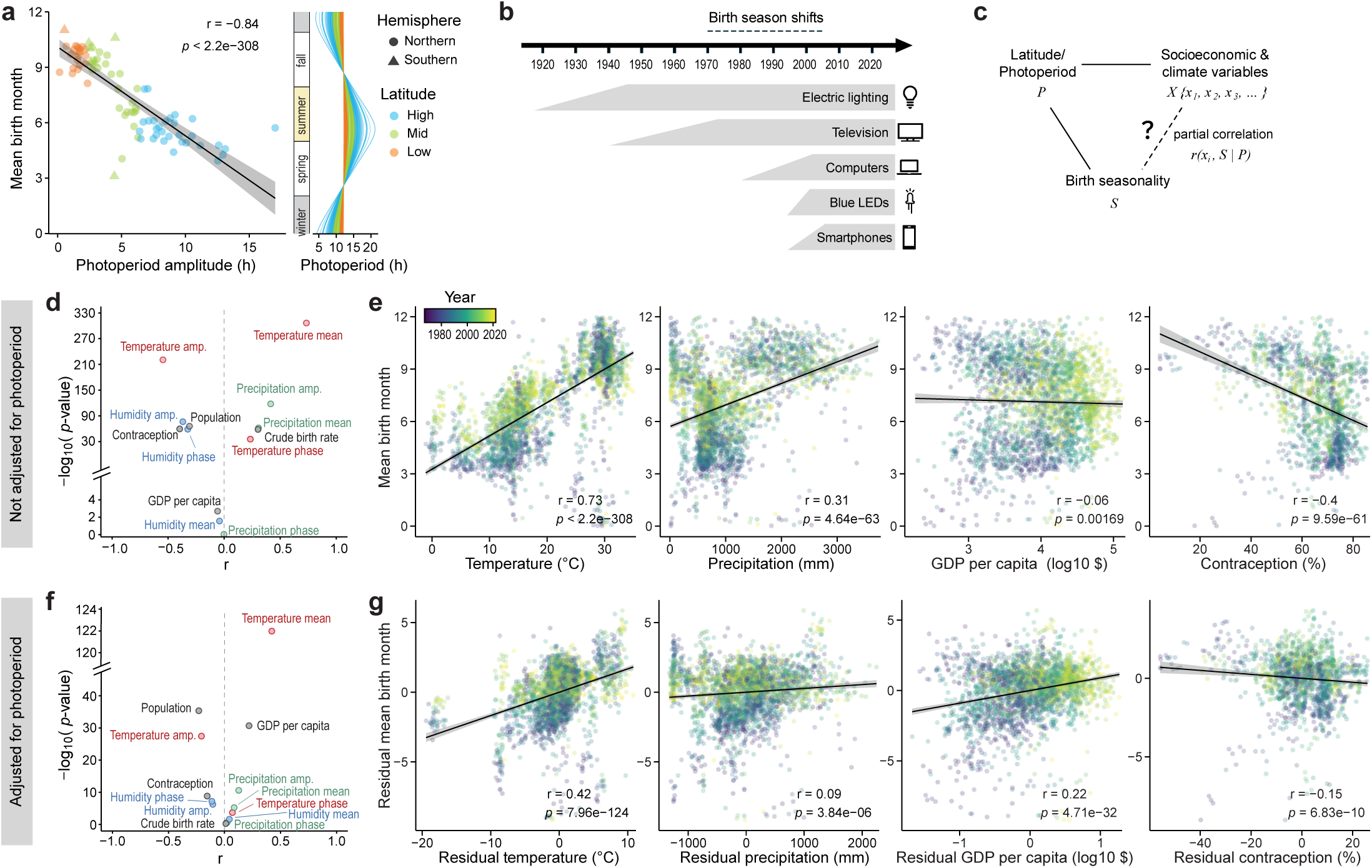
Temperature is the predominant predictor of birth peak timing. **a**, Scatter plot and linear regression of annual photoperiod amplitude versus birth peak phase (mean birth month) across countries/regions. The adjacent panel shows the annual photoperiod cycles for different countries/regions. **b**, Timeline of commercialization and widespread adoption of major technological innovations that could alter human light exposure. Note none of these technologies coincides with the onset and stabilization of the phase shift in birth peaks. **c**, Schematic of the correlation analysis used to test socioeconomic and climate predictors of birth peak timing, including partial correlations that adjust for potential covariation with latitude and photoperiod amplitude. **d, f**, Volcano plots showing Spearman correlation coefficients (r) versus statistical significance for each predictor, for both the original correlations (d) and partial correlations adjusting for latitude/photoperiod (f). **e, g**, Scatter plots and linear regression of example predictors versus birth peak phase across countries/regions and years, with (g) or without (e) adjustment for covariation with latitude/photoperiod. See also Fig. S5.

We therefore asked whether the observed birth peak phase delay might be due to technological innovations that alter light exposure in humans. To address this question, we examined timelines of initial commercialization and widespread adoption of key technologies in the developed world, including electric lighting, television, computers, blue LEDs, and smartphones. If any of these technologies had been a direct driver, birth peaks should have shifted shortly after their introduction and moved toward stabilization following widespread adoption. However, none of these timelines aligned well with the period of the rapid phase shift in birth peaks (Fig. 5b). Electric lighting and blue LEDs were of particular interest because they directly interfere with natural light-dark cycles and circadian entrainment.^48,49^ Yet, access to electric lighting began around the 1920s and became widely adopted around the 1940s in developed regions such as Europe,^50^ well before the onset of the birth peak shift. Blue LEDs were commercialized in the 1990s and became widely available in the 2000s,^51^ by which time the birth-peak phase shift was nearing stabilization. Together, these findings suggest that photoperiod and related technological changes are unlikely to drive the phase delay in birth peaks in recent decades.

### Temperature shows the strongest association with birth-peak timing

To identify other potential drivers underlying the observed variations in birth seasonality, we performed an unbiased analysis of 13 socioeconomic and climatic variables (Table S2). Socioeconomic variables included GDP per capita, contraceptive use, crude birth rate, and population size, with annual country/region-level data obtained from the World Bank and UN Data Portal.^33,52,53^ Climate variables included temperature, precipitation, and humidity, each represented as monthly time series for every country/region.^54^ From these data, we calculated annual mean, amplitude, and peak phase, resulting in a total of nine climate variables. Next, we examined correlations between each variable and birth peak timing across different years and regions. To account for covariation with latitude, we further used partial correlations controlling for photoperiod amplitude (Fig. 5c).

Surprisingly, this analysis revealed annual mean temperature as the predominant predictor of birth peak timing (Fig. 5d-g, Fig. S5b-c). Temperature was strongly correlated with birth peak phase (r = 0.73, *p* < 2.2e−308, Fig. 5d-e), comparable to that of photoperiod amplitude (r = −072, *p* < 2.2e−308, Fig. S5a). The correlation remained significant after adjusting for photoperiod (r = 0.42, *p* = 7.96e−124, Fig. 5f-g), suggesting that temperature could explain considerable variations in birth peak timing beyond those attributable to photoperiod or its covariation with latitude. In contrast, other variables showed much lower predictive power, with *p*-values more than 80 orders of magnitude weaker than that of mean temperature (Fig. 5f). For instance, GDP per capita was nearly uncorrelated with birth peak phase (r = −0.06, *p* = 0.00169, Fig. 5d-e). Although contraceptive use and mean precipitation were initially correlated with birth peak phase (contraception: r = −0.40, *p* = 9.59e−61; precipitation: r = 0.31, *p* = 4.64e−63; Fig. 5d-e), these correlations diminished after adjusting for photoperiod (contraception: r = 0.09, *p* = 3.84e−06; precipitation: r = −0.15, *p* = 6.83e−10; Fig. 5f-g). These findings are striking given the known influence of socioeconomic factors and human interventions on reproduction, as well as the role of precipitation in shaping seasonal climate patterns that exert strong evolutionary selective pressures in seasonal rhythms through food and water availability.

Assessing temporal trends of mean temperature change affirmed the findings above. Rate of temperature change was positively correlated with the rate of phase shift in birth peak timing (r = 0.46, Fig. 6a), indicating countries/regions with faster warming also showed faster phase delays of birth peaks. This agreed well with the observation that Europe, the fastest-warming continent after the Arctic,^55^ also had the largest phase delay of birth peaks.

**Figure 6.**
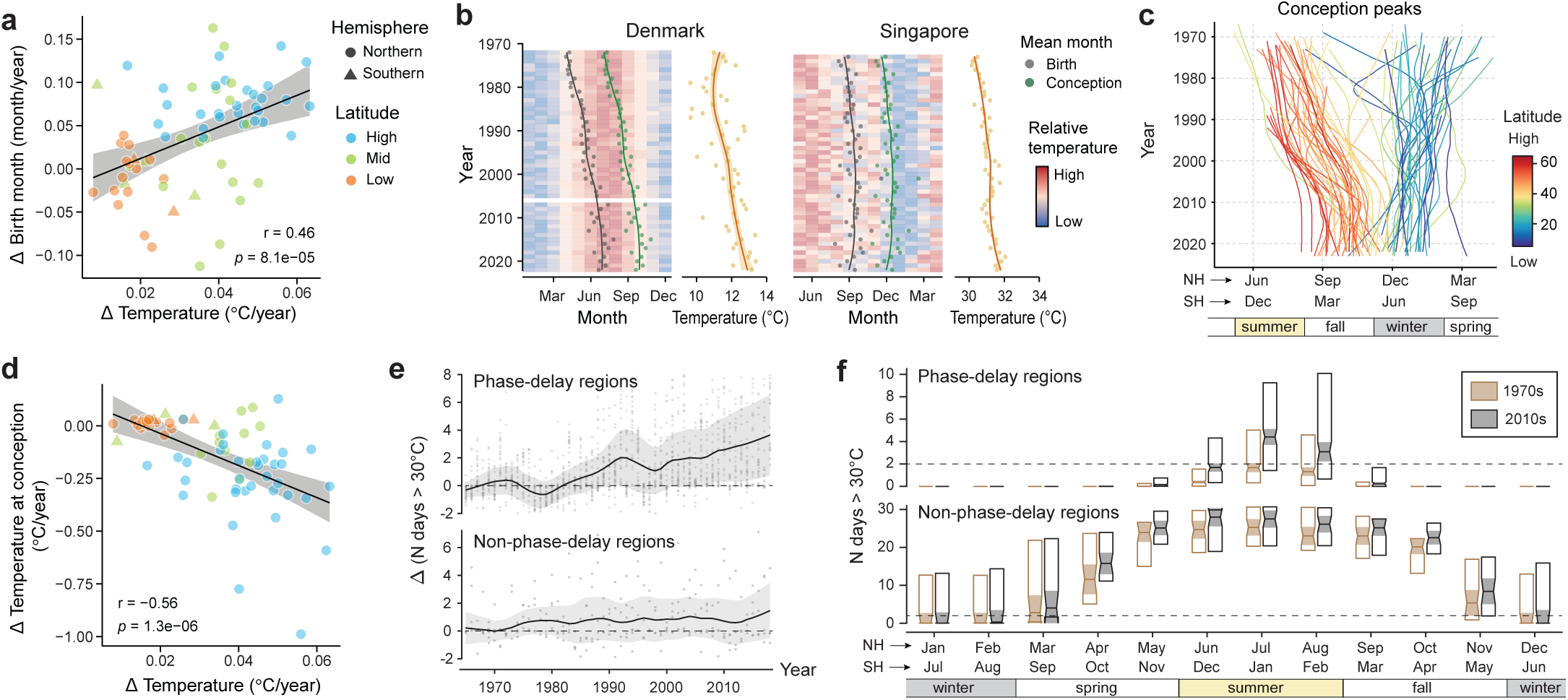
Conception peaks have shifted toward cooler months in a warming climate. **a**, Scatter plot and linear regression of the rate of change in annual mean temperature versus the rate of change in birth peaks across countries/regions. The rate of change was obtained by linear fitting (see Methods). A positive correlation indicates that faster warming regions tend to show faster phase delay in birth peak timing. **b**, Birth peak phase (gray) across years and its regression over time, and the inferred conception peak phase (green), estimated by shifting the birth peak nine months earlier. These are overlaid on a background heatmap showing relative monthly temperatures across years, shown separately for an example high latitude country (Denmark, left) and a low latitude country (Singapore, right). Adjacent panels show annual mean temperatures and their regressions (yellow) over the same time span for the two countries. **c**, Regression of yearly conception peak phases over time overlaid across different countries/regions (curves) and colored by absolute latitude, inferred based on birth peak data as in Fig. 4c. **d**, Scatter plot and linear regression of the rate of change in annual mean temperature versus the rate of change in temperature at the time of conception. Note the improved correlation coefficient in (d) compared with (a). **e**, Changes in the number of high-temperature days (Tmax>30°C) since the early 1970s in regions with (top) or without (bottom) a phase delay in birth peaks. Shown here are the averages of the two hottest months of the year (i.e., July and August in the Northern Hemisphere, January and February in the Southern Hemisphere). Black curves and shaded areas indicate LOWESS-smoothed means and standard deviations, respectively. **f**, Number of high temperature days (Tmax>30°C) across months of the year at the onset of birth peak phase delay (1970-1979) versus after its stabilization (2010-2018), shown separately for regions with (top) or without (bottom) a phase delay. Boxes show the median (center), 25th percentiles (bottom), and 75th percentiles (top), with notches and shading indicating the 95% confidence intervals of the median.

### Conception peaks shifted toward cooler months in a warming climate

Why did birth peaks shift toward hotter months in a warming climate? One possible explanation is that this shift reflects changes in conception timing. Moving birth peaks 9 months earlier revealed that conception peaks shifted toward colder months in high latitude regions (Fig. 6b-c). At lower latitudes, temperatures also rose, though at slower rates; however, conception already peaked during the coolest months of the year, leaving little room for further shifts. Indeed, a negative correlation (r = −0.56) was observed between the rate of mean temperature change and the rate of temperature change at the time of peak conception (Fig. 6d), supporting the hypothesis that conception peaks shifted toward cooler months in response to the warming climate.

One apparent caveat of this hypothesis lies in the disparity between the magnitude of climate warming and the much larger seasonal temperature differences involved (Fig. 6d). From the early 1970s to the 2000s, mean annual temperature increased by about 1-2°C at high latitudes. In contrast, the temperature at peak conception may have dropped by roughly 10°C, shifting from mid-summer to early/mid-fall. How could such modest long-term warming lead to such a large, non-linear response in seasonal timing?

A plausible explanation involves the rising frequency of high-temperature days and heatwaves. Despite only modest increases in annual mean temperature, the number of high temperature days (daily T_max_ >30°C) has grown substantially, with a steady upward trend in recent decades (Fig. 6e). Countries/regions with a phase delay in birth peaks showed a faster rise in high temperature days, with an average increase of 3.6 days per summer month over four decades, whereas those without a phase delay showed an average increase of less than 1.5 days during the same period.

Heat exposure is known to have detrimental effects on reproduction, and spermatogenesis is particularly heat sensitive.^56^ Heat shock to the testes can result in dysfunction and apoptosis of sperms and male germ/progenitor cells, reducing sperm count and quality.^56,57^ Although spermatogenesis is continuous and asynchronized, recovery after a single episode of heat stress may take weeks to a month.^56,57^ Thus, when the frequency of heatwaves surpasses a threshold, male reproductive function may remain at a suboptimal level throughout the hot summer months; in fall and winter, when heatwaves become much less frequent than summer, normal sperm production and quality may resume.

Consistent with this hypothesis, regions with a phase delay in birth peaks, typically at high latitudes, experienced a marked increase in summer heat exposure (Fig. 6f). In these regions, summer months generally had zero to two high temperature days in the 1970s, but this rose to an average of 2 to 5 days by the 2010s, coinciding with the shift in conception peaks to September or later, when high temperature days dropped to nearly zero (Fig. 6f). In contrast, regions without a phase delay, typically at mid-to-low latitudes, were already too hot during summer, and the number of high temperature days fell below two only in winter (Fig. 6f), consistent with their winter conception peaks.

Taken together, these results support the idea that increasing summer heat exposure may delay the optimal window for conception. This provides a biologically plausible mechanism by which conception peaks shift toward cooler months at high latitudes, even under modest warming of mean annual temperatures.

## Discussion

### Widespread seasonality in human births and deaths

Despite being classically viewed as non-seasonal, humans exhibit remarkably robust seasonal patterns in both births and deaths. Mortality peaks in mid-winter, and births peak from spring to fall depending on latitude, with both varying by ∼10-40% between peaks and troughs (Fig. 1-2, Fig. S1, Fig. S4). Our global analysis showed that these patterns are not confined to particular cultures, climates, or socioeconomic conditions, but instead represent widespread and persistent seasonal rhythms at the population scale. The long-term persistence of these cycles is striking. Even major societal disruptions such as the two World Wars caused only temporary perturbations, with birth seasonality rapidly returning to patterns similar to those before the war (Fig. 4e).

The global analysis also suggests that holiday-related conceptions do not dominate birth seasonality. Although the New Year holidays occur at roughly the same time worldwide, birth peaks did not always fall in September across regions (Fig. 2a-c). Southern hemisphere countries provide particularly compelling evidence. The minor September birth peaks in the mid-latitudes align with conceptions during New Year holidays, but are substantially weaker than the primary seasonal cycle that peaks around February and March (Fig. 2d). The timing of other major holidays also shows no correlation with birth peak timing (Fig. S1d).

### Latitudinal patterns in birth seasonality help reconcile regional inconsistencies

Reported patterns of human birth seasonality have long appeared inconsistent across regions. As early as 1835, Quetelet noted that “physiologists had already observed the influence of the seasons on human births and deaths; but in general, their results did not agree”.^10^ More than 150 years later, Roenneberg and Aschoff conducted an extensive analysis of human birth seasonality and similarly identified clear seasonal rhythms accompanied by substantial geographic variation in peak timing and waveform.^11,12^

By analyzing long-term demographic data from 124 regions spanning all major inhabited latitudinal zones, our results reveal that much of this apparent heterogeneity reflects a coherent latitudinal gradient in birth timing, with peaks occurring in spring to summer at higher latitudes and progressively transitioning to late summer and fall at lower latitudes in both hemispheres (Fig. 2a-c, Fig. S1c). Low latitude and tropical regions exhibit pronounced birth seasonality, often with greater amplitudes than those at higher latitudes (Fig. 2a-c). In contrast, deaths peak consistently in mid-winter across regions, except near the tropics where seasonality diminishes sharply (Fig. 2e-g, Fig. S1c).

Notably, the global analysis allowed a coherent pattern to emerge, which may reconcile long-standing puzzles of regional inconsistencies in birth seasonality. For example, it has long been unclear why the US exhibits summer-fall birth peaks, whereas Europe generally shows spring peaks.^11,12,21^ These differences align with the observed latitudinal gradient in birth timing, with the US mainly spanning mid-latitudes and Europe lying in higher latitudes. Previous work also reported puzzling bimodal birth patterns,^11,12,21^ which may result from holiday-related conception peaks that fall out of phase with the main seasonal cycle, and/or from mixing data across regions or years with different latitude or temperature distributions.

### Shifts in human reproductive timing linked to climate warming and increasing heatwaves

Systematic analysis of temporal trends, using data extending back to the 1840s, revealed consistent phase delays in birth seasonality across high latitudes over recent decades, whereas death patterns remained stable (Fig. 4, Fig. S4). Across all analyzed high-latitude regions, birth peaks shifted progressively from spring and early summer toward summer and fall, showing phase delays of around 2-3 months between the 1970s and early 2000s, followed by stabilization thereafter. This is consistent with a recent report by Hearn and Whitmore, that identified a similar phase shift in the UK around the mid-1970s.^24^ Our global dataset revealed that these phase delays were widespread, occurring across much of Europe and parts of North and South America, while most mid-low-latitude regions showed little to no change.

This phase shift does not appear to be explained by changes in photoperiod or by technological changes that alter human light exposure (Fig. 5a-b), pointing to potential non-photoperiod regulators of birth peak timing. The convergence of higher-latitude toward mid-low-latitude patterns suggests that a key underlying factor at high latitudes has shifted toward low-latitude conditions, and/or that birth timing reflects combined effects of multiple interacting factors.

An unbiased analysis of 13 socioeconomic and climatic variables identified temperature as the predominant predictor of birth-peak timing (Fig. 5c-g, Fig. S5). Warmer regions and years exhibited later peaks, even after controlling for latitude/photoperiod (Fig. 5e, g). Consistent with this finding, regions undergoing faster warming, such as Europe, also showed greater phase delays (Fig. 6a). This temperature association may also help explain why Chile, despite spanning mainly mid-latitudes, exhibited birth peak patterns resembling those of higher latitudes (Fig. 4a-b), as its population is concentrated in cooler, higher elevation areas. Hearn and Whitmore proposed an alternative hypothesis, that the introduction of freely available hormonal contraceptives might underlie the shift in birth timing in the UK.^24^ The global analysis presented here found only weak correlations between contraceptive use and birth peak timing, especially after controlling for latitude/photoperiod (Fig. 5d-g). Moreover, contraceptive use could not explain why high-latitude regions converged toward mid-low-latitude birth timing, nor why high-latitude regions, after shifting, stabilized at different phases along the summer-to-fall seasonal spectrum in a manner that follows their latitude (Fig. 4c).

While it initially seemed counterintuitive that birth peaks shifted toward hotter months in a warming climate, analysis of estimated conception timing revealed that conception peaks shifted toward cooler months at high latitudes (Fig. 6b-d). In contrast, conception at mid-low latitudes already peaked in winter, leaving little room for further shifts despite rising temperatures (Fig. 6b-c). This pattern aligns with the well-established heat sensitivity of male reproductive biology.^56^ Spermatogenesis is negatively impacted by high temperatures, and recovery can take weeks to a month.^57^ Thus, the increasing frequency of summer heatwaves, even amid modest changes in annual mean temperature, could compromise male fertility during summer months and push conception toward cooler times of the year. Indeed, high latitude regions with phase delays experienced marked increases in summer heatwaves, from around 0-2 high-temperature days/month in the 1970s to 2-5 days/month in the 2010s, and the delayed conception peaks coincide with the seasonal decline in high-temperature days (Fig. 6e-f). Consistent with this hypothesis, mid- and low-latitude regions were already too hot in summer, leaving winter as the most favorable window for conception (Fig. 6f). This interpretation aligns well with existing evidence on human sperm parameters. Meta-analyses have reported higher sperm count and quality in winter than in summer,^58,59^ and a broad literature has documented long-term declines in sperm count and quality since the 1970s.^60–62^

Together, these data suggest that climate change, especially the rising frequency of heatwaves at high latitudes, contribute to the shift in seasonal timing of human births. This introduced a new dimension of climate-health interactions. Increased heatwaves may contribute to the declining male fertility,^60–62^ in addition to their well-recognized role in driving heat-related mortality^29–32^ (Fig. 2h). Changes in population birth timing may also have broader epidemiological implications, given known associations between season of birth and the incidence of various childhood and adult diseases.^14,15^ These findings underscore the importance of understanding the biological basis of seasonal variations in human health and the broader demographic consequences of climate-driven changes in human seasonality.

### What is the biological basis of human seasonality?

Although temperature could explain both spatial and temporal variations in birth seasonality, the biological basis of seasonality in human births and deaths remains unknown. Their divergent latitudinal and temporal trends suggest that birth and death seasonality are governed by partially separate mechanisms rather than a single unified pathway.

Winter death seasonality provides interesting insight into the physiological relevance of these rhythms. Winter mortality strengthens consistently with age across multiple countries (Fig. 3a-c, Fig. S2), indicating that winter mortality disproportionately affects older populations. Consistent with previous reports,^18,19,34–41^ cause-specific analyses showed that this pattern is driven by a broad set of diseases, including cardiovascular, respiratory, neurological, digestive, endocrine, and infectious conditions, but not neoplasms or external causes (Fig. 3d, Fig. S3). Analysis of influenza surveillance data further revealed that shifts in flu seasons, such as the unusually early 2009-2010 flu season, did not advance the timing of winter mortality peaks (Fig. 3f-h). Together, these findings suggest a generalized seasonal vulnerability that amplifies with age and involves multiple organ systems, rather than a single biological process or isolated behavioral or epidemiological trigger.

More broadly, the global presence of seasonality in human births, deaths, and disease vulnerability suggests that the human body undergoes coordinated seasonal changes. This idea is supported by a growing body of molecular and clinical evidence. Previous studies have reported seasonal patterns in hormones, gene expression, circulating cytokines and peptides, adipose tissues, brain functions, and sleep.^25,63–71^ The convergence of demographic, clinical, and molecular evidence suggests a multi-organ, body-wide seasonal modulation spanning immune functions, endocrine signaling, metabolic control, and brain activity.

Despite these advances, the origins and mechanisms of human seasonality remain poorly understood. Although molecular mechanisms underlying circannual rhythms remain unknown even in animals,^1,47^ the coherence and persistence of human seasonal patterns raise the possibility that humans may also possess a form of circannual control, potentially acting alongside multiple environmentally entrained pathways. These may include ambient temperature, photoperiod, food availability, and other modifying environmental factors, which interact synergistically to entrain seasonal physiological timing. This work showcases the value of systematic, population-level approaches for studying biological timing in real-world human populations.^25,63,72–74^ The findings here define an empirical foundation for future mechanistic studies of human seasonality and highlight the need for interdisciplinary work at the intersection of chronobiology, physiology, genetics, epidemiology, and climate science.

## Methods

### Birth, death, latitude, and flu data collection

Monthly counts of births and deaths were obtained from the United Nations Database (UNdata),^33^ which compiles vital statistics reported by national/regional statistical offices, civil registration systems, and population censuses. UNdata harmonizes these submissions, resolves inconsistencies where possible, and standardizes country-month-year records into a unified format. These raw datasets included a total of 159 countries/regions (156 with birth records, and 140 with death records). Birth data range between 1967-2024, while death data range between 1980-2024.

We also obtained longer-term monthly birth data of selected countries using the Human Fertility Database (HFD).^46^ In addition, we obtained mortality data by age, sex, and/or underlying causes of death for selected countries directly from individual national statistical offices, including Switzerland,^75^ New Zealand,^76^ Netherlands,^77^ UK,^78^ and US.^42^ Flu data were obtained from World Health Organization’s Global Influenza Programme FluID dataset.^43^

For each country/region’s latitude, we manually compiled the most populous city and its corresponding latitude and longitude coordinates using Google search.

### Preprocessing and filtering of birth, death, and flu data

Birth and death records from UNdata were first preprocessed to ensure consistency across regions and years. For each country/region, all records were first grouped and deduplicated by prioritizing entries labeled “Final figure, complete” in the UN reliability field; when multiple observations existed for the same year-month combination, only the most reliable entry was retained. We also removed non-standard entries, such as 3-month aggregated entries. To ensure complete temporal coverage, only years with all 12 months present were kept. To ensure robustness of the analysis, we filtered countries/regions with insufficient counts. Filtering was conducted separately for births and deaths. Specifically, a region was retained only if it had at least three years of data with annual means ≥300, or at least eight years with annual means ≥100. To ensure robustness, we also applied the thresholding across years in downstream temporal analyses. The final filtered dataset included a total of 124 countries/regions, with birth records from 122 countries/regions spanning 1967-2023 and death records from 95 countries/regions spanning 1980-2023, covering regions across six continents. For death and flu data periodicity analysis, we excluded data from 2020 and beyond, to avoid the confounding effects of the COVID-19 pandemic. This also resulted in the removal of two additional countries from the death analysis due to their insufficient numbers of remaining years, producing a final total of 93 countries/regions for the death periodogram analysis.

### Quantification of seasonal cycle significance, amplitude, and peak phase

Because individual months differ in length, raw monthly counts were first adjusted by the number of days in each month, including accounting for the February difference in leap years. Data were then normalized within each year by dividing by the annual sum of the adjusted counts, yielding adjusted monthly fractions. Next, we computed monthly deviations from a uniform expectation of 1/12 by subtracting this baseline from each month’s adjusted fraction, dividing the difference by 1/12, and converting the result to a percentage. This resulted in monthly percent deviations that indicate the extent to which monthly birth rates are higher or lower relative to a flat, nonseasonal distribution. Schematic and formulas of these preprocessing steps were also shown in Fig. 1d.

To detect periodicities, we applied Lomb-Scargle periodograms for each time-series using the Matlab function plomb (spectrumtype set to ’normalized’). Significance (*p* values) was estimated using the Baluev method.^79^

Averaged seasonal cycles of birth and death were obtained for each country/region by calculating the median of monthly deviation across all years. Average seasonal cycle amplitude was then calculated as half the difference between the maximum and minimum of these monthly medians. We applied two arbitrary thresholds to categorize countries/regions into high (> 43°), mid (>20° and ≤ 43°), and low (≤ 20°) latitude zones. The average seasonal cycle for each latitude zone was calculated as the median of monthly deviation across all countries/regions within the latitude zone.

We next calculated seasonal cycle amplitude and peak phase for each region and year, for regions that are highly seasonal with a 12-month periodicity significance of *p* < 1e-10. The yearly amplitude was defined as the half difference between the maximum and minimum monthly values. We used circular statistics to calculate peak phase. Specifically, monthly values were converted to angular positions on the unit circle (2*π*·(month–1)/12) and weighted by their normalized monthly fractions to obtain an angular mean month. This angular mean month was then converted back to calendar-month space to represent the estimated peak phase. To obtain a more robust estimate of peak phase, we computed the center of mass of the weighted monthly deviations across all 12 months. We also evaluated the center of mass computed from the top 1 to top 6 months, which yielded similar but noisier results. To calculate the average seasonal cycle peak phase for each country/region, we also used circular statistics and averaged peak phase across all years with data. To align peak phase by season between the two hemispheres, all southern hemisphere data (latitude <0) were phase shifted by 6 months. Note that all numerical values of peak phase (i.e., mean birth month) shown in the figures (Fig. 5-6, Fig. S5) use such hemisphere-aligned values, with [0 1] corresponding to January in the Northern Hemisphere and July in the Southern Hemisphere. Flu and climate variable peak timing was calculated similarly using circular statistics as the birth peak phase.

To analyze temporal trends of seasonal cycle amplitude and peak phase across years, we further restricted the analysis to countries/regions with at least 20 years of data in the UN database. Trends were assessed using both linear regression and nonparametric LOWESS smoothing. Circular statistics were applied when analyzing peak phase. Analyses were performed either by grouping regions into high-latitude versus mid-low-latitude categories (Fig. S4, Fig. 4d) or by examining individual regions separately (Fig. 4c). The leftmost curve in Fig. 4c corresponds to South Korea, and is driven by a disproportionate concentration of birth registrations in January and February. This pattern likely reflects a known reporting artifact associated with systematic biases in birth registration timing^80^ and was therefore excluded from subsequent analyses.

### Analysis of socioeconomic and climate variables

To search for potential extrinsic regulators of birth peak phase, we assembled a set of socioeconomic and climatic predictors, including photoperiod, major holiday, GDP per capita, population size, crude birth rate, contraceptive prevalence, temperature, precipitation, and humidity.

For photoperiod, we first calculated the annual photoperiod cycle for each country/region based on its latitude. Daylength for a given latitude *ϕ* and day of year *d* was computed using the following formula,^81^

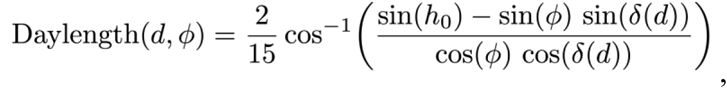

where h₀ = –0.833°, representing the solar altitude at sunrise/sunset, and

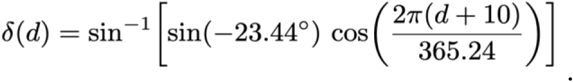

Annual photoperiod amplitude was then defined as the absolute difference between daylength on the summer solstice and the winter solstice.

Date or calendar month of major holidays for all analyzed countries/regions were manually compiled using Google search.

GDP (1960-2023, 262 countries/territories) and population size (1960-2023, 265 countries/territories) were obtained from the World Bank’s World Development Indicators (WDI) dataset,^52^ which compiles national statistics from annual government reporting systems. GDP per capita was calculated as total GDP divided by population of the corresponding country/region. Both GDP per capita and population size were then log10 transformed.

Crude birth rate was calculated from UNdata’s monthly birth counts and population size of the corresponding country/region. For each country/region and year, annual total births were divided by population size of the matched country/region and year to obtain an annualized crude birth rate.

Contraceptive prevalence was obtained from the United Nations’ World Contraceptive Use dataset (1970-2023; 258 countries/territories).^53^ Following standard UN recommendations, we restricted analyses to estimates corresponding to “Married or in-union women” derived using the UN’s interpolation-based methodology to maximize comparability across countries and years.

Climatic data (temperature, precipitation, and humidity) were obtained from the World Bank Climate Change Knowledge Portal, which provides country-aggregated monthly climate statistics derived from the CRU TS v4.07 dataset (0.5° × 0.5° gridded coverage, 1901-2022, monthly resolution, ∼237 countries and territories).^54^ For temperature, the dataset contains monthly averaged values for T_max_, T_min_, and T_avg_ (daily maximum, minimum, and average temperatures), which are highly correlated. We found similar results across all three indicators, and reported T_max_ here to avoid redundancy. For each climate factor, we calculated three variables for downstream analysis, including the annual mean, cycle amplitude, and peak phase for each country/region and year. Amplitude was defined as the difference between the maximum and minimum monthly value. Peak phase (mean month of the annual cycle) was estimated using circular statistics as described above, including alignment of the phase of Northern and Southern Hemispheres by season.

We next calculated Spearman’s rank correlations between each predictor and birth peak phase at the region-year level, both with and without adjusting for latitude/photoperiod. For each predictor, we first calculated a simple bivariate correlation with birth peak phase (i.e., without adjustment of latitude/photoperiod). Note that data coverage differed across predictors. Correlations were calculated using only country-year combinations with complete data for each predictor pair. To account for potential covariance with latitude/photoperiod, we additionally calculated partial correlations by controlling for latitude (photoperiod amplitude). This was done by performing two separate linear regressions: 1) between the predictor and photoperiod and 2) between birth peak phase and photoperiod. Residues were extracted from each regression, and Spearman’s correlation was then computed between the two sets of residues to obtain the partial correlation. Each correlation analysis provided a coefficient and an associated *p*-value. Multiple testing across predictors was performed using the Benjamini-Hochberg false-discovery rate correction.

### Conception timing and heatwave analysis

Temporal trends in seasonal birth timing were estimated for countries with ≥20 years of birth data and ≥15 years of temperature data. For each country, annual mean birth months were first computed using circular statistics as described above. These annual means were then represented as angular values on the unit circle, and the sine and cosine of these angular values were each regressed linearly against year. The resulting regression coefficients were used to directly estimate the average angular rate of change of the mean birth month over time, and were subsequently converted from radians per year to units of months per year, yielding an estimate of the direction and magnitude of temporal shifts in mean birth timing. Countries were classified as phase-delay regions if their temporal slope of mean birth month was > 0.035 month/year. The mean conception month for each country-year was then estimated as nine months before the observed mean birth month.

Rates of change in annual mean temperature (°C/year) were calculated as the slope of a linear regression of T_max_ against year for each country. T_max_ was from the CRU TS v4.07 dataset (see above). Next, we estimated the rate of temperature change at the time of conception for each country/region. Because conception peak phase can be fractional values, we linearly interpolated the temperature at the conception peak phase for each year using the two neighboring months of the same year. We then performed linear regression of interpolated temperatures across all years for each country/region, where the fitted slope represents the rate of temperature change at the time of conception.

High-temperature exposure was quantified using the HadEX3 climate extremes dataset,^82^ a global gridded product produced by the WMO Expert Team on Climate Change Detection and Indices. HadEX3 provides monthly indices of climate extremes from 1901-2018 on a 1.25° latitude × 1.875° longitude grid, derived from quality-controlled station observations interpolated to a consistent global field. For this analysis, we used the TXge30 index, which represents the number of days with daily maximum temperature (T_max_) exceeding 30 °C, providing a standardized measure of high temperature days. To obtain country-level estimates, each country/region was assigned the value from the grid cell containing the coordinates of its most populous city. Txge30 data for Southern Hemisphere countries/regions were phase shifted 6 months after to align the data by season. For the temporal trend analysis, we limit the analysis to 1965 and onward due to significant (>5%) lack of data from many regions before the time.

We then calculated the change in the number of high temperature days for the two hottest summer months (i.e., July and August for Northern Hemisphere; January and February for Southern Hemisphere) since the early 1970s separately for phase-delay regions and non-phase delay regions. Phase-delay regions were defined as those with a birth month shift slope of >0.035 month/year (i.e., ∼1 month over three decades), and vice versa for non-phase delay regions. We next subtracted raw TXge30 entries with the median Txge30 of the first 10 years (i.e., 1965-1974), and performed LOWESS smoothing over aggregated data of all countries/regions under the category. We also analyzed Txge30 for each month, comparing the first 10 years at the onset of birth peak phase delay (1970-1979) versus after its stabilization in the 2010s (2010-2018).

### Data and Materials Availability

Monthly birth and death data used in this study were obtained from the United Nations Database (UNdata) (https://data.un.org/). Long-term monthly birth data for selected countries were obtained from the Human Fertility Database (https://www.humanfertility.org/Data/DataAvailabilityShort). Additional birth and mortality data were obtained from national statistical offices, including the United States Centers for Disease Control and Prevention WONDER underlying cause-of-death database (https://wonder.cdc.gov/), Statistics New Zealand (https://www.stats.govt.nz/information-releases/births-and-deaths-year-ended-march-2025/), the Swiss Federal Statistical Office (https://www.bfs.admin.ch/), the UK national statistical authority (https://www.ons.gov.uk/peoplepopulationandcommunity/birthsdeathsandmarriages/deaths), and Statistics Netherlands (https://www.cbs.nl/en-gb/figures/detail/70895ENG). Influenza data were obtained from the World Health Organization Global Influenza Programme FluID dataset (https://www.who.int/teams/global-influenza-programme/surveillance-and-monitoring/influenza-surveillance-outputs). Contraceptive prevalence data were obtained from the United Nations World Contraceptive Use database (https://population.un.org/dataportal/home). Climate data were obtained from the World Bank Climate Change Knowledge Portal (https://climateknowledgeportal.worldbank.org/download-data). High-temperature exposure data (TXge30) were obtained from the HadEX3 climate extremes dataset (https://www.metoffice.gov.uk/hadobs/hadex3/download_etsci.html). Gross domestic product (GDP) and population size data were obtained from the World Bank World Development Indicators (https://data.worldbank.org/indicator/NY.GDP.MKTP.CD; https://data.worldbank.org/indicator/SP.POP.TOTL). Holiday timing data and country latitude coordinates were manually compiled through Google search. Custom code and metadata from the study will be deposited in a public repository.

## Acknowledgements

We are grateful to many colleagues for their insightful discussions at various stages of the study, particularly Drs. Joseph Takahashi, Shin Yamazaki, Carla Green, William Dauer, Zhengda Li, Daniel Jarosz, Mark Krasnow, Fabienne Aujard, Martine Perret, and Jacques Epelbaum.

## Author Contributions

Conceptualization: S.L.

Methodology: S.L., C.L., N.S., J.E.F.;

Software: C.L., S.L.

Validation: C.L., N.S., S.L.;

Formal Analysis: S.L., N.S., C.L.;

Investigation: S.L., N.S., C.L.;

Resources: S.L., J.E.F.;

Data Curation: N.S., C.L., S.L.;

Writing -original draft: S.L., C.L., N.S.

Writing -review & editing: S.L., J.E.F.

Visualization: S.L., C.L.;

Supervision: S.L.;

Project Administration: S.L.

Funding Acquisition: J.E.F., S.L.

## Supplementary tables

**Table S1.** Analyzed countries/regions with corresponding birth and death seasonality parameters.

**Table S2.** Socioeconomic and climate variables analyzed and their summary statistics.

---

**Figure S1.**
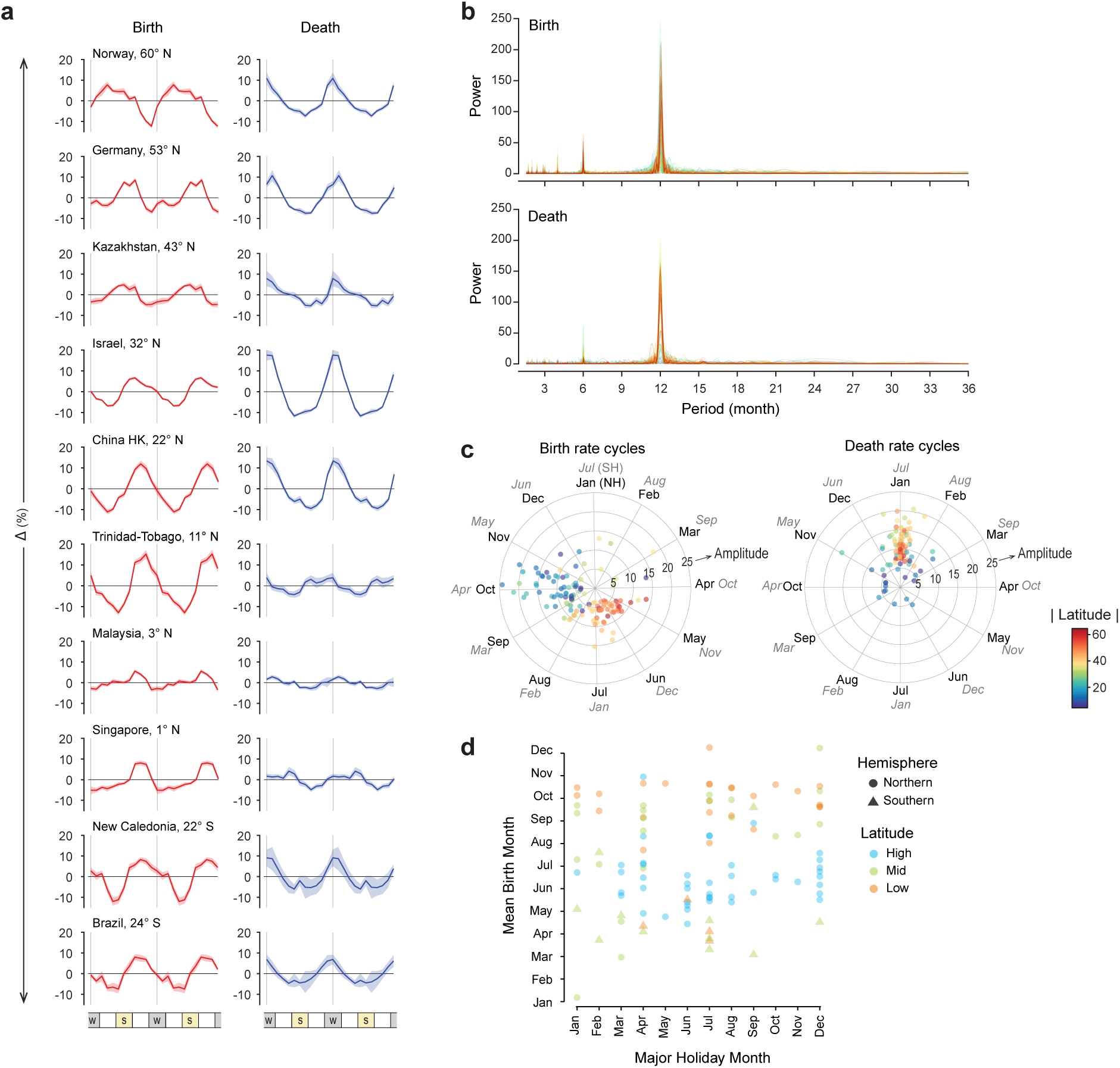
Global patterns of seasonality of births and deaths in humans. **a**, Average annual patterns and 95% confidence intervals of Δbirths (left) and Δdeaths (right) in example countries/regions, averaged across years. Patterns are double plotted for two cycles for visualization purposes. Vertical gray lines indicate mid-winter (i.e., January in the Northern Hemisphere and July in the Southern Hemisphere). **b**, LS periodograms of monthly birth (top) and death (bottom) rates across countries/regions detected significant 12-month periodicity. **c**, Polar plots showing birth (left) and death (right) peak phase and amplitude averaged across years for each country/region. Countries/regions (curves in b, dots in c) are colored by absolute latitude. Inner month labels denote Northern Hemisphere (NH); Outer month labels denote Southern Hemisphere (SH). **d**, Scatter plots of major holiday months versus mean birth peak months across countries/regions.

**Figure S2.**
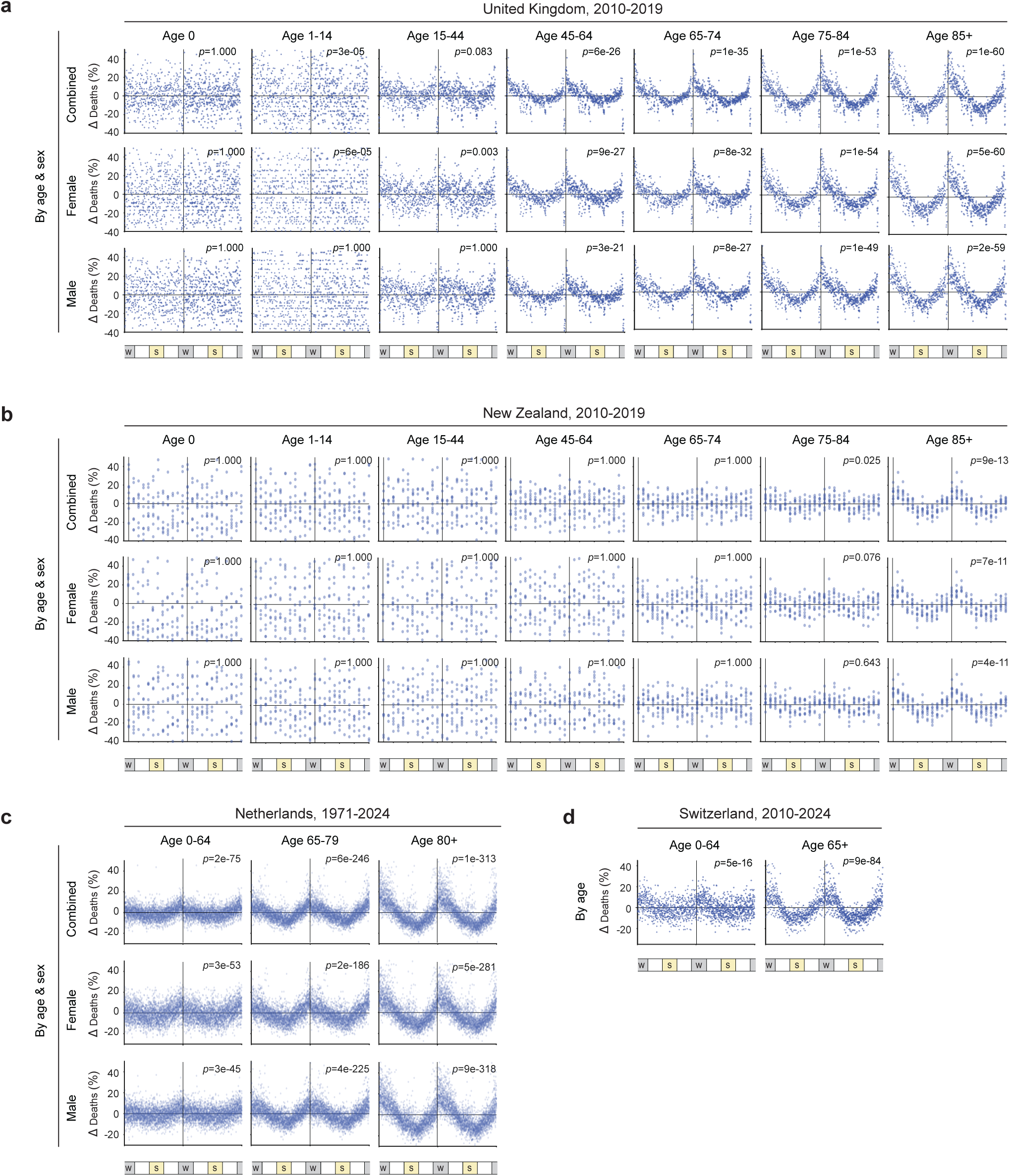
Death seasonality strengthens with age. **a-d**, Δdeaths in the UK (a), New Zealand (b), the Netherlands (c), and Switzerland (d) across different age groups and/or sexes, as reported by national statistical agencies. Panels of Δdeaths are formatted similarly as in Fig. 3c-d. Note the strengthening of death seasonality with increasing age across all countries.

**Figure S3.**
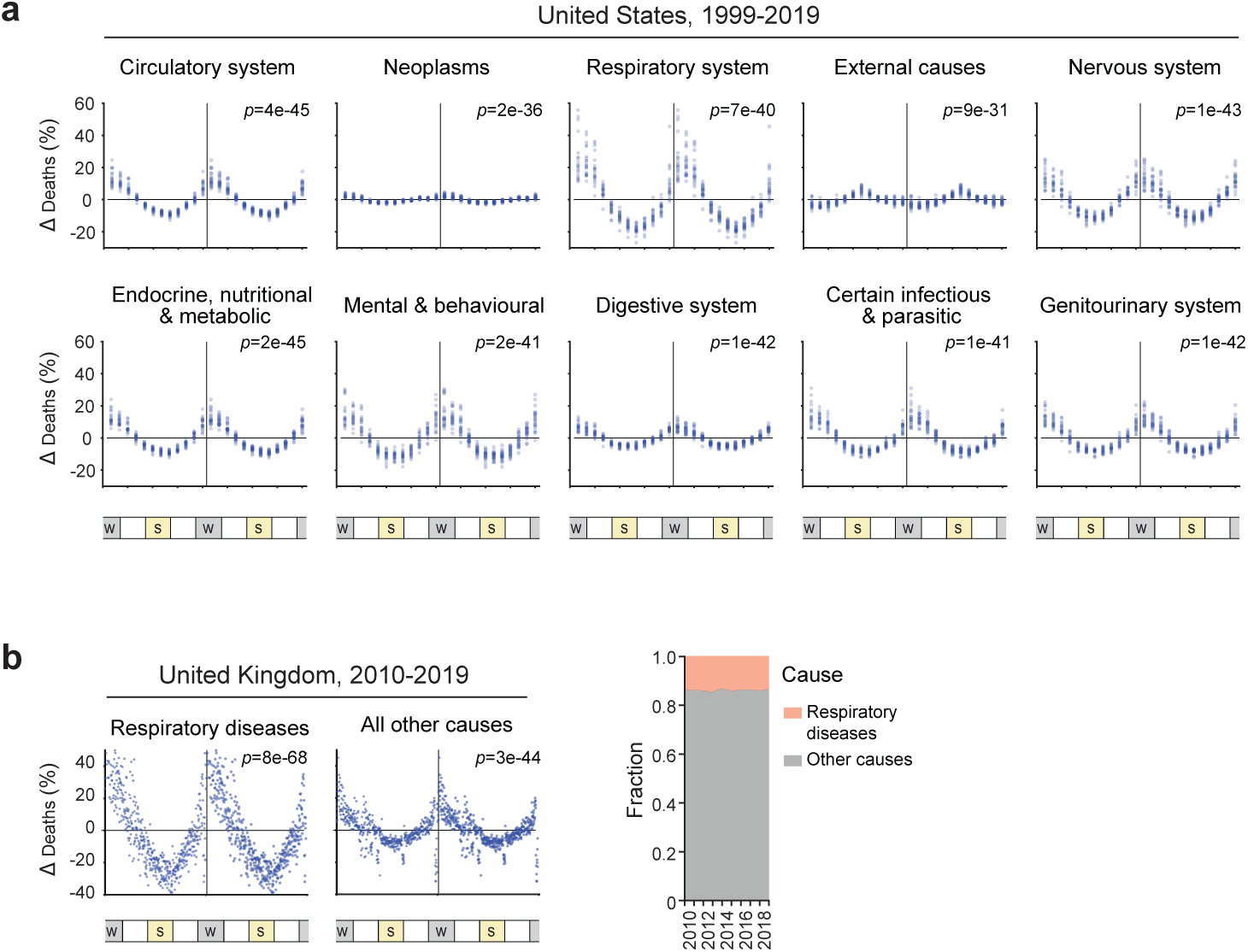
Death seasonality by cause of death. **a**, Δdeaths in the US for the top 10 leading causes of death, shown as scatter plots. Data are of monthly resolution. **b**, Δdeaths (left) and fractions of death (right) in the UK for respiratory diseases versus non-respiratory causes, shown as scatter plots. Data are of weekly resolution. Panels of Δdeaths are formatted similarly as in Fig. 3c-d.

**Figure S4.**
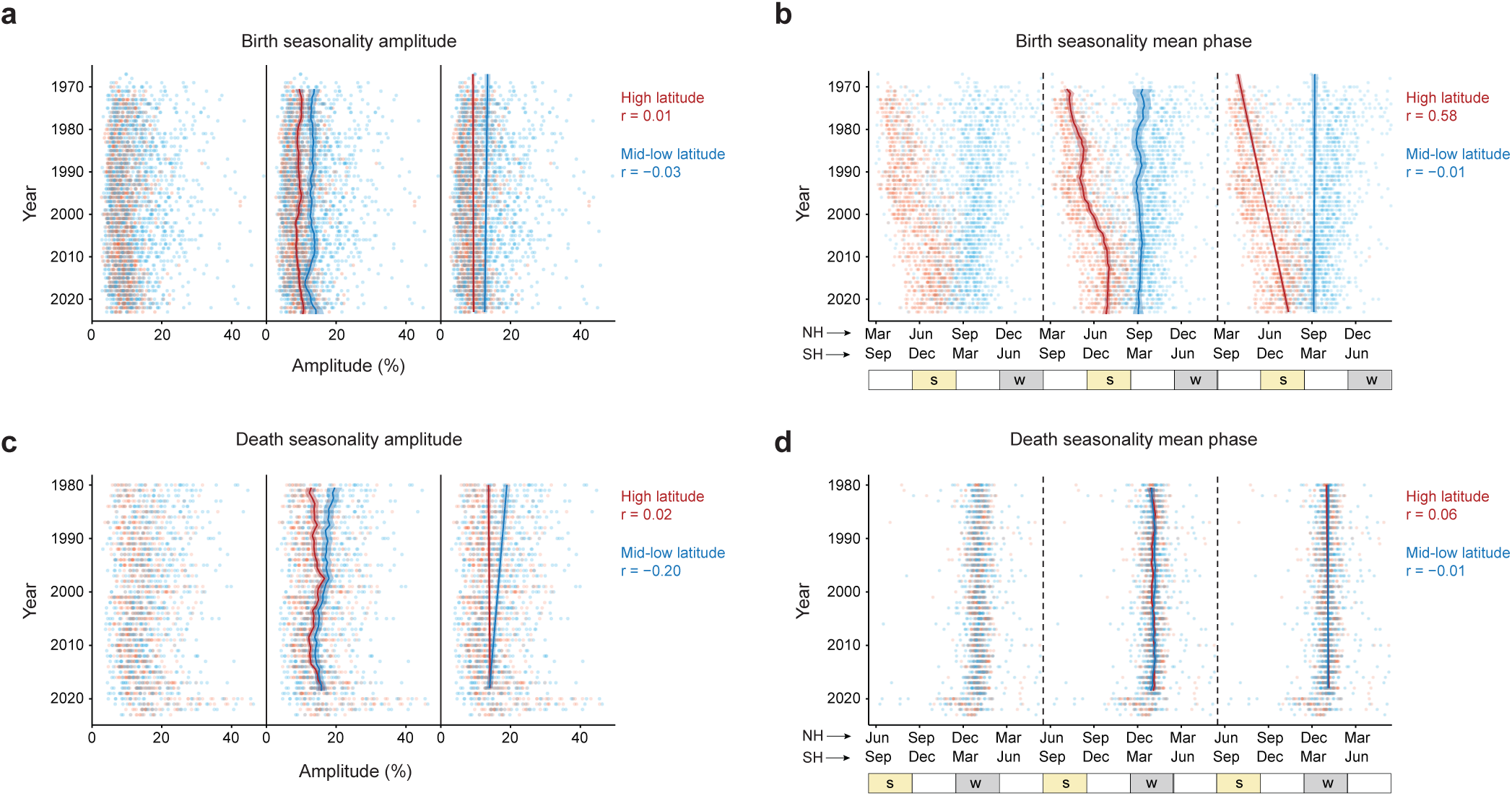
Temporal patterns in the amplitude and peak phase of birth and death seasonality. **a-d**, Seasonal cycle amplitude (a, c) and peak phase (b, d) for individual country/region and year (dots), shown separately for birth (a, b) and death (c, d) data. Within each panel, data are triple-plotted to demonstrate the distribution of individual data points (left), non-parametric LOWESS smoothing (middle), and linear regression (right) over time for high-and low-latitude regions.

**Figure S5.**
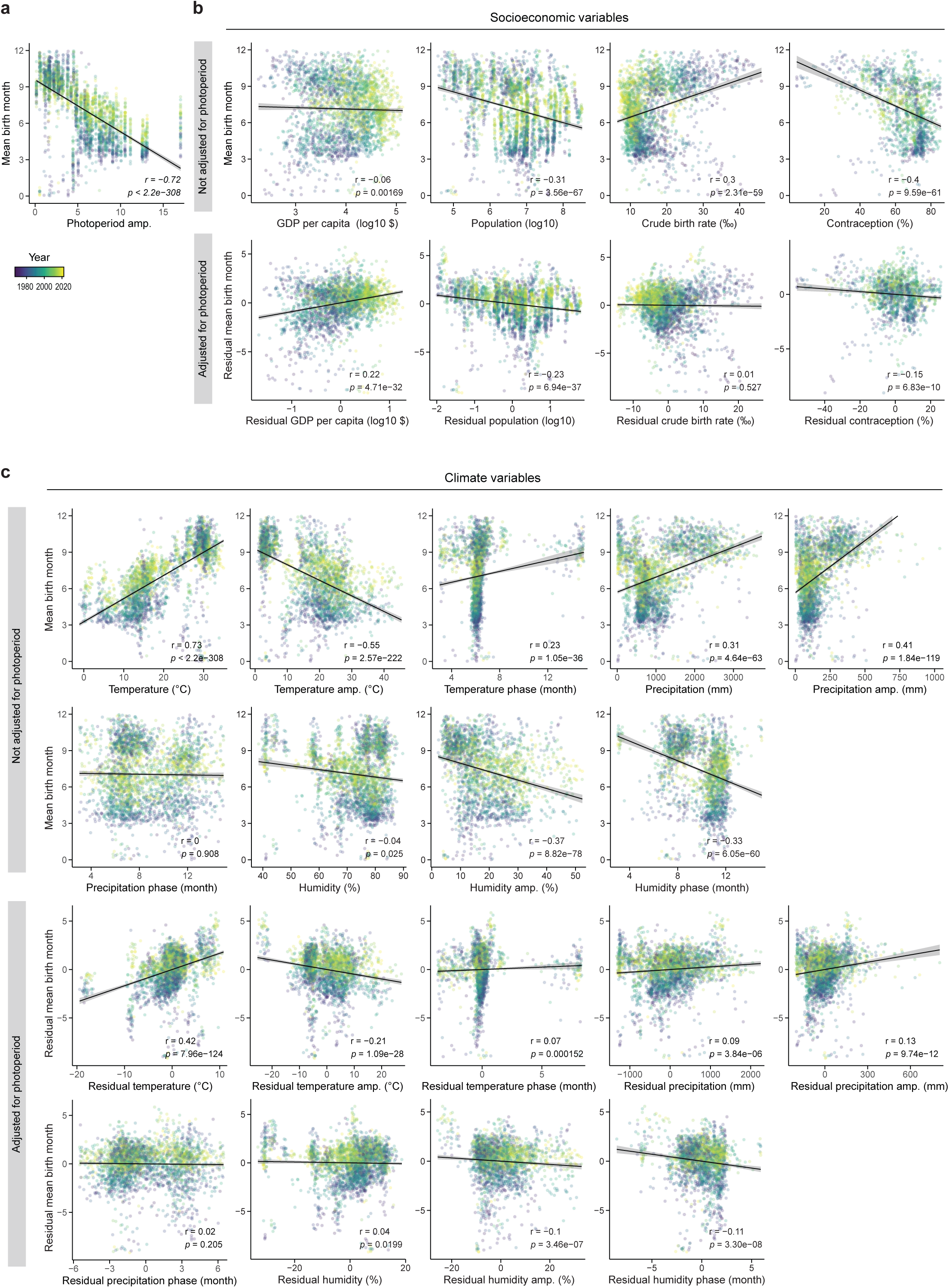
Correlations of 13 socioeconomic and climatic variables with temporal and geographic variation in birth peak phase. **a**, Scatter plots and linear regression of photoperiod amplitude versus birth peak phase across countries/regions and years. **b-c**, Scatter plots showing each socioeconomic (b) and climate (c) predictors versus birth peak phase across countries/regions and years, with (bottom) or without (top) adjustment for covariation with latitude/photoperiod.

